# Cold acclimation has a differential effect on leaf vascular bundle structure and carbon export rates in natural Arabidopsis accessions originating from southern and northern Europe

**DOI:** 10.1101/2020.04.14.040428

**Authors:** Katja Schneider, Lorena Abazaj, Cornelia Niemann, Laura Schröder, Thomas Nägele

## Abstract

Acclimation to low but non-freezing temperature represents an ecologically important process for *Arabidopsis thaliana* but also for many other plant species from temperate regions. Cold acclimation comprises and affects numerous molecular and physiological processes and the maintenance of sugar supply of sink tissue by photosynthetically active source tissue is preliminary for plant survival. Here, we analysed the correlation of changes in vascular bundle structure at the leaf petiole and sucrose exudation rates before and after cold acclimation. We compared six natural Arabidopsis accessions originating from southern and northern Europe. Photosynthetic capacities, i.e. maximum and effective quantum yield of photosystem II, revealed a significant effect of condition but not of genotype. Only for northern accessions we observed a highly significant negative correlation between leaf sucrose exudation rates, xylem and petiole cross section areas. Further, only for northern accessions we observed a significant increase of vascular bundle and leaf petiole cross section area during cold acclimation. In contrast, variance of cross section areas of cold acclimated southern accessions was strongly reduced compared to control plants while mean areas remained similar under both conditions. In summary, our findings suggest that natural Arabidopsis accessions from northern Europe significantly adjust sink strength and leaf vascular bundle structure to stabilize plant growth and photosynthesis for survival under low and freezing temperature.

## Introduction

The exposure of higher plants to low but non-freezing temperature induces a tightly regulated, multigenic and multifaceted process termed cold acclimation. Cold acclimation is a crucial process for many plant species of the temperate zone because low temperature significantly impacts on their range boundaries (Hoffmann, 2002). It comprises and affects gene expression, translational and post-translational processes and induces significant alteration in metabolic pathway regulation (Wang *et al.*, 2017, Bahrani *et al.*, 2019, Liu *et al.*, 2019). To sustain a stable metabolic homeostasis during cold exposure, photosynthetic capacities and metabolism of carbohydrates need to be tightly adjusted to prevent toxic accumulation of reactive oxygen species (ROS) (Dreyer and Dietz, 2018). Although it is known that plants alter the composition and activation state of their photosynthetic apparatus due to environmental changes within a process termed photosynthetic acclimation, many of the involved molecular mechanisms are still not understood. Regulation of metabolic pathways, and particularly of sucrose (Suc) metabolism, are well known to play a central role in photosynthetic acclimation (Strand *et al.*, 2003, Stitt *et al.*, 2010, Nägele *et al.*, 2012, Herrmann *et al.*, 2019). In the light, photosynthetic CO2 fixation results in triose phosphates which are substrate for starch biosynthesis in the chloroplast and Suc biosynthesis in the cytosol which is regulated by cytosolic fructose-1,6-bisphosphatase (cFBPase) and sucrose phosphate synthase (SPS) (Strand *et al.*, 2000). Suc represents a central transport sugar in many plant species, it plays an important role as storage compound, represents an osmotically active solute and is involved in sugar signalling (Ruan, 2014). Recent findings suggest that Suc export from the chloroplast into the cytosol is involved in cold acclimation and is required for development of maximal freezing tolerance (Patzke *et al.*, 2019). The authors discussed that Suc in the chloroplast serves as a reservoir to supply cytosolic and vacuolar sugar metabolism under stress conditions. Supportingly, our own studies indicated that Suc compartmentation and invertase-driven cleavage in cytosol and vacuole significantly affect cold stress response and stabilization of photosynthesis (Nägele and Heyer, 2013, Weiszmann *et al.*, 2018).

In addition to its role in metabolic regulation and subcellular compartmentation, Suc is the main transport sugar in *Arabidopsis thaliana* and many other plant species. Stabilization of Suc supply for carbon sink organs, e.g. roots, by photosynthetically active source tissue is essential for plant growth and development under changing environmental conditions. For transport, Suc is loaded into the phloem which ultimately establishes a source-sink hydrostatic pressure differential which drives phloem transport. In the sinks, Suc is unloaded and provided for cellular metabolism. Readers interested in details of phloem loading, transport and unloading, please refer to more detailed and specialized literature at this point (e.g. (Johannes and Patrick, 2017)). In general, Suc is secreted into the cell wall space by passive SUAGRS WILL EVENTUALLY BE EXPORTED TRANSPORTERs (SWEETs) and loaded into the phloem by SUCROSE TRANSPORTERs (SUCs or SUTs) (Braun, 2012). A recent study demonstrated that sucrose transporter SUC2, which regulates carbon export from leaves of *Arabidopsis thaliana,* is controlled via ubiquitination and phosphorylation which directly links whole plant carbon homeostasis to posttranslational modification (Xu *et al.*, 2020). Further, previous studies have shown that regulation of SUTs is part of abiotic stress response (see e.g. (Gong *et al.*, 2015)) and that both transcriptional and posttranscriptional mechanisms are involved in regulation of Suc export (Xu *et al.*, 2018).

Due to the availability of natural accessions with a broad longitudinal and latitudinal range of geographical origin, *Arabidopsis thaliana* has become an attractive system to study plant-environment interactions, ecological and evolutionary patterns (Bouchabke *et al.*, 2008, Mishra *et al.*, 2014, Mönchgesang *et al.*, 2016, The 1001 Genomes Consortium, 2016, de Jong *et al.*, 2019). Cold acclimation capacity differs significantly between natural Arabidopsis accessions, and accessions with differential cold and freezing tolerance significantly differ in regulation of photosynthesis and carbohydrate metabolism (Hannah *et al.*, 2006, Fürtauer *et al.*, 2019, Zuther *et al.*, 2019). Yet, many aspects still remain unclear about the stabilization of carbohydrate homeostasis under low temperature. Particularly, natural variation of regulatory and developmental processes involved in the stabilization sink-source interactions remain elusive. In addition to the molecular level, e.g. the regulation of SUC2, also structural and anatomical properties, e.g. sieve elements (SE) and companion cells (CC) of minor veins (Davidson *et al.*, 2011), need to be addressed.

In the present study, we analysed natural variation of leaf Suc exudation and vascular bundle structure at leaf petioles during cold acclimation. We compared cross section areas of vascular bundle tissue and recorded leaf Suc exudation rates under 22°C and 4°C in six natural accessions originating from southern and northern Europe. Additionally, we recorded chlorophyll fluorescence parameters to monitor photosynthetic capacities before and after cold acclimation.

## Materials and Methods

### Plant material and growth conditions

Plants of natural *Arabidopsis thaliana* accessions C24 (southern accession, Portugal), Ct-1 (southern accession, Italy), Fei-0 (southern accession, Portugal), Col-0 (northern accession, Germany), Oy-0 (northern accession, Norway) and Rsch-4 (northern accession, Russia) used in this study were grown on a 1:1 mixture of GS90 soil and vermiculite in a climate chamber under short-day conditions (8h/16h light/dark; 90 μmol m^-2^sec^-1^; 22°C/16°C; 70% relative humidity). After 4 weeks, plants were transferred to the greenhouse and grown under long day conditions (16h/8h light/dark). After 7 days in the greenhouse, plants were either (i) sampled (microscopy, exudation) and used for chlorophyll fluorescence measurements (“control”), or (ii) transferred to a cold room for temperature acclimation (4°C, 16h/8h light/dark; 90 μmol m^-2^sec^-1^). After 7 days at 4°C, cold acclimated plants were sampled (microscopy, exudation) and used for chlorophyll fluorescence measurements.

### Chlorophyll fluorescence measurements

The maximum quantum yield of PSII (Fv/Fm) was determined after 15 min of dark adaptation by supplying a saturating light pulse. Dynamics of quantum efficiency of PSII (ΦPSII), photochemical (qP) and non-photochemical quenching (qN) were determined within light response curves by stepwise increase of photosynthetically active radiation (PAR) from 0 to 1200 μmol m^-2^ s^-1^ in 5 min intervals. All measurements were performed using a WALZ^®^ GFS-3000FL system equipped with measurement head 3010-S and Arabidopsis chamber 3010-A (Heinz Walz GmbH, Effeltrich, Germany, https://www.walz.com/). Cold exposed plants were acclimated to ambient temperature, i.e. 22°C, for 20 minutes before measurement.

### Sample preparation, staining and semi thin sectioning for light microscopy analysis

Petioles of *Arabidopsis* plants were cut into 1 mm pieces in buffer (75 mM cacodylate, 2 mM MgCl_2_, pH 7.0) and stained over night at room temperature in a 1:1 mixture of safranin/astra blue (1% (w/v) safranin in ddH_2_O/ 0,1% (w/v) astra blue, 2% (w/v) tartaric acid). Samples were dehydrated with ethanol (2x 10 min 50% (v/v) EtOH; 3x 100% EtOH for 10, 30 and 60 min) and embedded in Spurr’s resin. For each genotype under both analysed conditions, at least 25 cuts from at least 5 different plants were prepared. For preparation of semi thin sections, 5 samples of each ‘genotype x condition pool’ were randomly chosen for further preparation. Semi thin sections (thickness, 2 μm) were cut using a diamond knife on a Reichert Ultracut-E ultra-microtome. Sections were mounted on glass slides and examined using a Zeiss Axiophot light microscope. Images were acquired using a SPOT insight 2 MP CCD colour digital camera. Light microscopy images were used to measure the areas of total petiole, vascular bundles and xylem. Area of other vascular bundle (VB) tissue than xylem was calculated as the difference of total VB area and xylem area. Examples for images and area definitions of petiole and VB tissues are provided in the supplements (Supplementary Figure S1 and S2).

### Sucrose quantification from exudates

To determine the sucrose concentration from exudates rosettes from plants grown under control conditions and after cold acclimation were placed in a petri dish containing 20 mM EDTA solution (pH 8.0). Leaves were cut with a razor blade at the centre of the rosettes. Five leaves from each plant were pooled and placed in a 1.5 ml reaction tube filled with 20 mM EDTA (pH 8.0) and exposed to growth PAR intensity, i.e. 90 μmol m^-2^sec^-1^, for 1h. Exudation was stopped by removing the leaves from the tube. Leaves were carefully dried on a paper towel and weighed on a fine scale to determine the fresh weight per exudate. The entire exudate was dried in a desiccator. Exudation experiments were performed at 22°C with control plants and at 4°C with cold acclimated plants. Dried exudates were resolved in 300μl ddH_2_O. 100μl of solved exudate was mixed with 100μl 30% (w/v) KOH and boiled at 95°C for 10 minutes. After quick cooling on ice 1ml of anthrone reagent (0.14% (w/v) anthrone in 14.6M sulfuric acid) was added, briefly mixed by inverting and incubated at 40°C for 30 minutes. Extinction was measured immediately at 620 nm.

### Statistics and area measurements

For statistical data evaluation we used the free software environment R Version 3.6.1 (https://www.r-project.org/) (R Core Team, 2019) and RStudio Version 1.2.5019 (https://rstudio.com/) (RStudio Team, 2019). Area measurements were done in ImageJ 1.52A using the ‘Area’ tool (https://imagej.nih.gov/ij/index.html).

## Results

### Geographical origin has no significant effect on photosynthetic capacities of cold acclimated plants

In all accessions, maximum quantum yield of PSII (Fv/Fm) significantly dropped from 0.82-0.83 under control conditions to 0.77-0.78 after cold acclimation (Fig. 1; ANOVA, p < 0.05). In accession Fei-0, Fv/Fm dropped to the lowest mean level (0.75) which was, however, not significantly lower than in other accessions. Fv/Fm was recorded following 15 minutes of leaf dark adaptation. Following Fv/Fm measurement, light response curves were recorded to determine dynamics of quantum efficiency of PSII (ΦPSII), qP and qN under different PAR intensities. Similar to Fv/Fm, also ΦPSII and qP were significantly reduced in cold acclimated plants compared to control plants across all accessions (ANOVA, p<0.001), while no significant ‘genotype effect’ was observed. For both parameters, the difference between mean values of control and cold acclimated plants increased with PAR intensity applied within the light response curves (Supplemental Figure S3, S4). A significant ‘genotype effect’ was observed for non-photochemical quenching (qN; ANOVA p <0.05) which was, in contrast to all other accessions, only weakly affected in the Portuguese accession C24 (Supplemental Figure S5). In summary, and with the exception of qN in C24, cold acclimation had a significant effect on all quantified photosynthetic parameters while no significant effect was found to differentiate accessions from southern and northern Europe.

**Figure 1.**
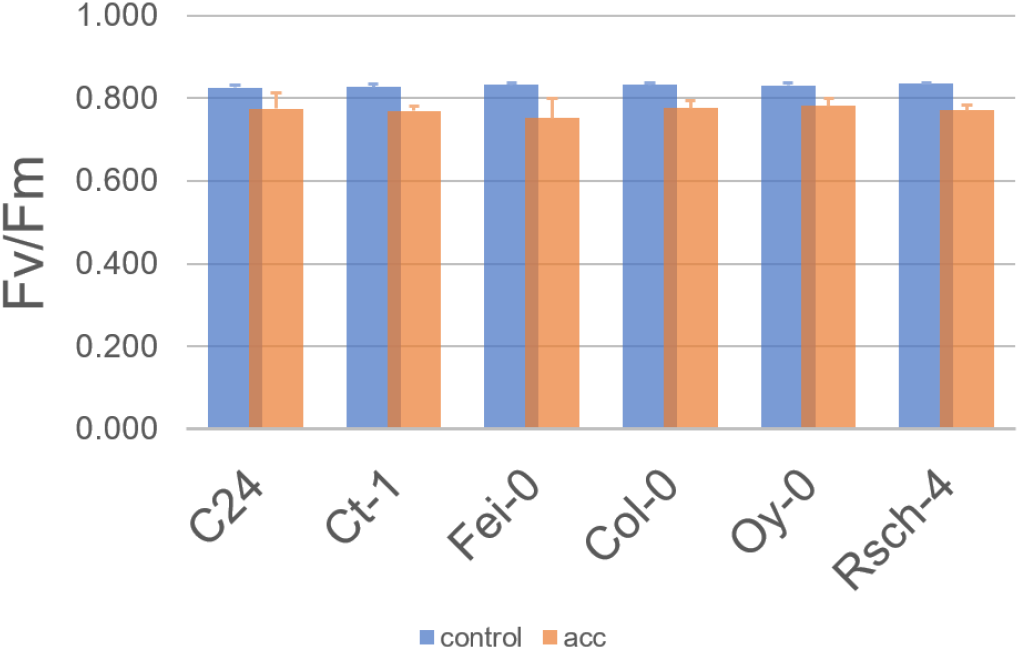
Maximum quantum yield of PSII in accessions before and after cold acclimation. Bars represent mean values +/− SD (n = 3). Blue bars: control plants; orange bars: cold acclimated plants (7d at 4°C).

### The reduction of leaf exudation rates of sucrose during cold acclimation is more distinct in accessions from northern than from southern Europe

The stabilization of carbon supply of non-photosynthetically active sink organs, e.g. roots, by source leaves is essential for successfully acclimation to low temperature. We quantified the exudation rate of sucrose for control plants and cold acclimated plants under respective growth temperature, i.e. at 22°C for control plants and at 4°C for cold acclimated plants. Under control conditions, northern accessions had a significantly higher leaf exudation rate than southern accessions (Fig. 2; ANOVA, p<0.001). In contrast, after 7d of cold acclimation, no significant difference between exudation rates quantified at 4°C was observed anymore between southern and northern accessions. Remarkably, exudation rates in two out of three southern accessions, C24 and Fei-0, dropped only slightly and not significantly at 4°C. Exudation rates in Ct-1 and all northern accessions decreased significantly during cold acclimation (ANOVA, p<0.05). The most distinct reduction of exudation rates was observed in the Russian accessions Rsch-4 from ~0.5 to 0.07 μmol C6 gFW^-1^ h^-1^.

**Figure 2.**
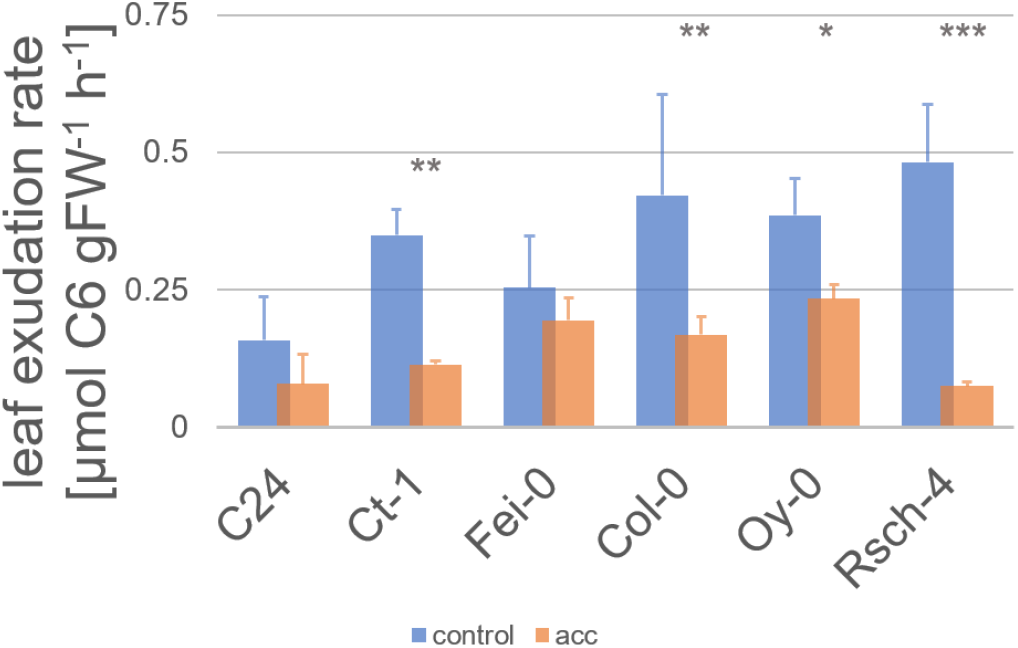
Leaf exudation rates before and after cold acclimation. Bars represent mean values +/− SD (n = 5). Blue bars: control plants; orange bars: cold acclimated plants (7d at 4°C). Asterisks indicate significance (ANOVA). * p<0.05; ** p<0.01; *** p<0.001.

### Cold acclimation induces structural changes in vascular bundle structure at the leaf petiole

Evaluation of xylem and other VB tissue areas from light microscopy images of leaf petiole cross sections revealed a positive correlation across all accessions and conditions (Fig. 3). Under control conditions, xylem and other VB areas were significantly correlated in southern accessions with R^2^ = 0.9924 and p<0.0001 (Pearson correlation). In contrast, only a weak positive (R^2^=0.0322) and non-significant correlation was observed in northern accessions. This changed during cold acclimation and resulted in a significantly positive correlation (R^2^=0.4998, p<0.01).

**Figure 3.**
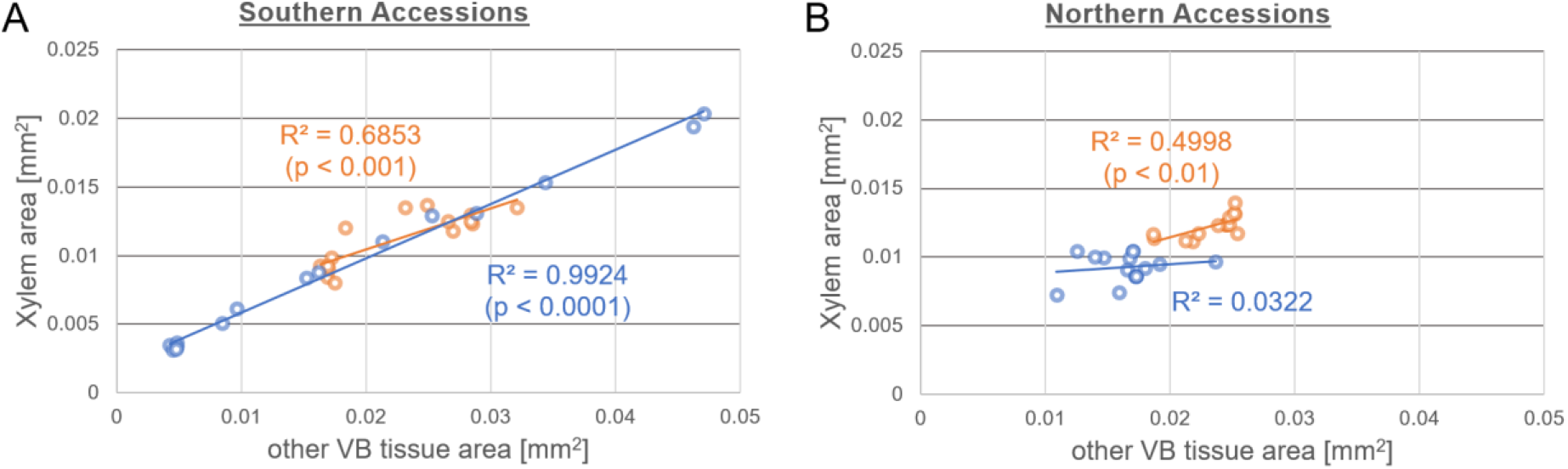
Correlation of xylem and other VB tissue area at leaf petioles. (**A**) Areas derived from microscopy of southern accessions C24, Ct-1 and Fei-0. (**B**) Areas derived from microscopy of northern accessions Col-0, Oy-0 and Rsch-4. Blue coloured: control samples; orange coloured: cold acclimated samples. Significance is indicated by p-values of Pearson correlations.

Also, in cold acclimated plants of southern accessions such a significant and positive correlation was observed (R^2^=0.6853, p<0.001), which, however, compared to control conditions, decreased both in regression coefficient and significance.

### Leaf exudation rate negatively correlates with xylem and petiole area in cold acclimated plants of northern accessions

Leaf exudation rates were correlated with cross section areas of xylem, other VB tissue and the complete petiole (Fig. 4). In the southern accessions C24, Ct-1 and Fei-0, cold acclimation induced a positive and nearly significant correlation of leaf exudation rate and xylem area (Fig. 4 A). Correlations with other VB tissue and petiole area remained similar to those observed under control conditions (Fig. 4 B, C). Interestingly, the variance of all areas in southern accessions became smaller due to cold acclimation. For example, xylem areas ranged between ~0.003 – 0.02 mm^2^ under control conditions but only between ~0.008 – 0.015 mm^2^ in cold acclimated plants (Fig. 4 A). A similar effect was observed for other VB tissue and petiole area (Fig. 4 B, C).

**Figure 4.**
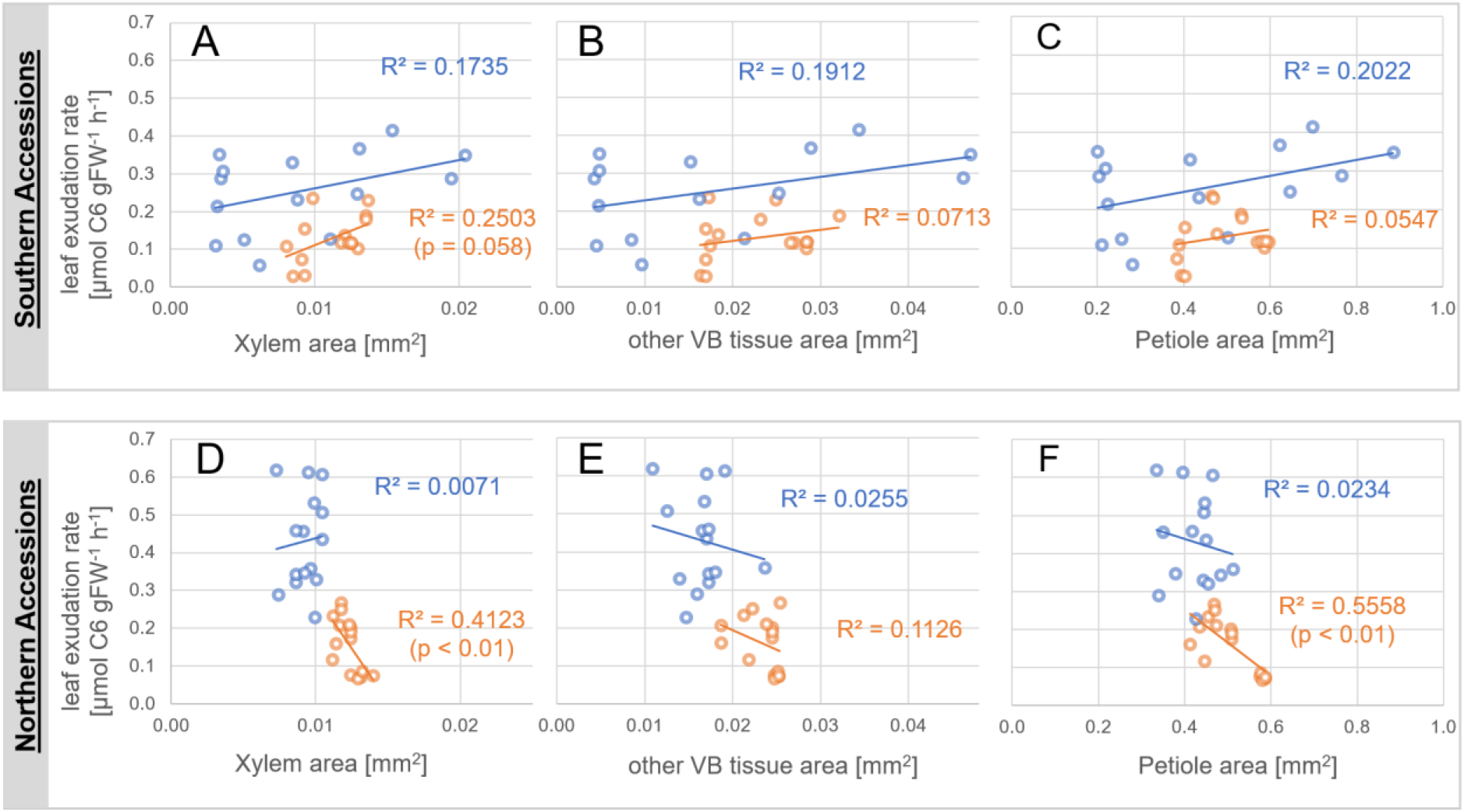
Leaf exudation rates as function of cross section areas. Upper panel (**A-C**): leaf exudation rate as function of (**A**) xylem area, (**B**) other VB tissue area, (**C**) petiole area of southern accessions C24, Ct-1 and Fei-0. Lower panel (**D-F**): leaf exudation rate as function of (**D**) xylem area, (**E**) other VB tissue area, (**F**) petiole area of northern accessions Col-0, Oy-0 and Rsch-4. Blue coloured: control samples; orange coloured: cold acclimated samples (7d, 4°C). Significance is indicated by p-values of Pearson correlations.

In northern accessions, i.e. Col-0, Oy-0 and Rsch-4, variance of all cross section areas remained similar before and after cold acclimation (Fig. 4 D-F). However, as already indicated in Figure 3, xylem areas significantly increased due to cold acclimation (p<0.001, ANOVA) from a mean area of 0.009 mm^2^ before to 0.0123 mm^2^ after cold acclimation (increase ~32%). Other VB tissue and petiole areas also increased significantly by ~41% and ~20%, respectively.

In contrast to southern accessions, cold acclimation induced a significant and negative correlation of leaf exudation rates with xylem areas as well as with petiole areas in northern accessions (Fig. 4 D, F; p<0.01). Although not significant, also other VB tissue area was negatively correlated to leaf exudation rates before and after cold acclimation. Thus, in summary, cold acclimation of northern accessions resulted in a pronounced and significant change of xylem areas which finally resulted in a negative correlation with leaf exudation rates.

### Statistical integration of structural and physiological elements reveals significance of cold acclimation output in accessions from northern Europe

Principal component analysis (PCA) coupled with Hotelling’s T^2^ statistics revealed a significant separation of control and cold acclimated samples of northern accessions within a 95% confidence region, while, in contrast, confidence ellipses of southern accessions overlapped (Fig. 5). In both southern and northern accessions photosynthetic parameters, i.e. Fv/Fm, ΦPSII, qP and qN similarly contributed to the separation of both conditions. Yet, while in southern accessions the exudation rate factor loading was oriented in nearby 90° angle towards xylem, other VB tissue and petiole areas (Fig. 5 A), the angle was almost 180° in northern accessions which strongly contributed to separation of both conditions along PC1 (Fig. 5B). Among southern accessions, acclimation response differed between the Italian accession Ct-1 and both Portuguese accessions C24 and Fei-0. Control and cold acclimated samples of Ct-1 were predominantly separated on PC2 which was determined by the significant effect on exudation rates (Fig. 5A and Fig. 2). In C24 and Fei-0, control and cold acclimated samples were separated on PC1 which was, predominantly, determined by photosynthetic acclimation. Northern accessions displayed a more homogenous pattern and cold acclimation was found to be almost exclusively explained by PC1 which explained 20% more of the total variance than in southern accessions (54% vs 74%; Fig. 5B). In summary, northern accessions displayed a conserved cold acclimation response in which photosynthetic acclimation was consistently associated with acclimation of leaf exudation rates and increased cross section areas at the leaf petiole. In southern accessions cold acclimation response was more diverse and less significantly associated with quantified cross section areas at the leaf petiole.

**Figure 5.**
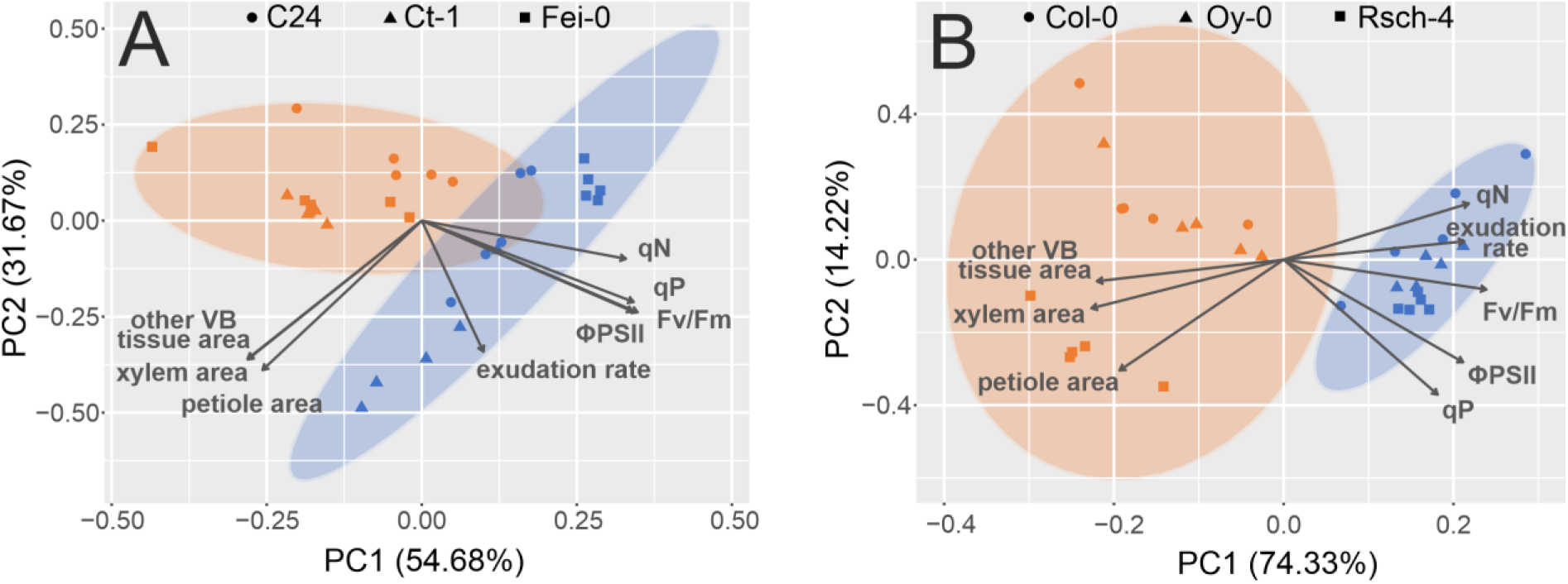
Principal component analysis of structural and physiological parameters. (**A**) PCA of southern accessions. Symbols represent independent samples of different genotypes: •C24, ▲ Ct-1, ■Fei-0. Blue coloured: control samples; orange coloured: cold acclimated samples (7d 4°C). (**B**) PCA of northern accessions. Symbols represent independent samples of different genotypes: •Col-0, ▲ Oy-0, ■Rsch-4. Blue coloured: control samples; orange coloured: cold acclimated samples (7d 4°C). Arrows represent loadings. Confidence regions (95%) are indicated by coloured ellipses. Data were scaled to zero mean and unit variance (autoscaling, z-scores).

## Discussion

Stabilizing carbohydrate supply of sink tissue by photosynthetically active source tissue is essential for plant acclimation to a changing environment. Particularly under low temperature, this stabilization is challenged by several factors. In source leaves, photosynthesis and carbohydrate metabolism need to be immediately adjusted if temperature drops to prevent a misbalance of photosynthetic primary and secondary reactions and irreversible cell and tissue damage. Although we observed slight differences between qN of accessions, our data indicated a similar capability of accessions from southern and northern Europe to photosynthetically acclimate to 4°C. This finding is supported by previous studies which did not detect a correlation between classical chlorophyll fluorescence parameters, i.e. Fv/Fm, ΦPSII or qP, and cold tolerance of natural Arabidopsis accessions at low but non-freezing temperatures (Ehlert and Hincha, 2008, Distelbarth *et al.*, 2013, Mishra *et al.*, 2014). Further, recorded light response curves indicated a similar capability of all accessions to cope with a dynamic range of PAR intensities before and after cold acclimation. Together with our previous studies, where we observed similar CO2 assimilation rates of cold acclimated plants of southern and northern accessions at 4°C (Nägele *et al.*, 2011, Nägele *et al.*, 2012, Nägele and Heyer, 2013), this provides further evidence for similar photosynthetic capacities of all accessions under these conditions. Discrimination of cold sensitive and tolerant accessions was shown to be possible using a different experimental and measurement design which applies combinatorial imaging together with the analysis of chlorophyll fluorescence transients of whole plant rosettes following mild sub-zero temperature treatments (Mishra *et al.*, 2014). In summary, these findings suggest that although cold acclimated plants of accessions originating from southern and northern Europe have similar photosynthetic capacities at 4°C, cold acclimation of northern accessions results in higher capacities to survive a temperature drop into the sub-zero range.

Accessions from southern and northern Europe differed significantly in their vascular bundle structure which was estimated from transversal sections at the leaf petiole. In northern accessions, cold acclimation induced a positive correlation between areas of xylem and other VB tissue, comprising also the phloem. In contrast, in southern accessions correlations were significant already under control conditions indicating less pronounced cold-induced developmental effects on VB tissue. Remarkably, total variance of VB tissue areas was reduced during cold acclimation of southern accessions which may suggest that VB tissue development is more constrained by low temperature than in northern accessions. Yet, also in context of light acclimation and phloem cell numbers of minor leaf veins, southern European accessions, here form Italy, were found to acclimate less pronounced than Swedish accessions (Adams *et al.*, 2018). Further, greater minor cross sectional sieve element areas found in high versus low light acclimated plants of Swedish accessions were hypothesised to be a prerequisite for their higher photosynthetic capacities (Cohu *et al*., 2013, Adams *et al*., 2018). While in our present and previous studies no significant genotype-effect was observed with regard to photosynthetic capacity under low temperature, in summary these findings suggest a generally less pronounced acclimation response in the development of vascular tissue under changing abiotic conditions and a clear limitation by low temperature in accessions from south Europe. Remarkably, however, this lowered acclimation response did not result in a lowered rate of leaf sucrose exudation compared to northern accessions at 4°C (see Fig. 2). Beyond that, two out of the three southern accessions rather seemed to stabilize exudation rates at 4°C to reach a value similar to that under control conditions, i.e. 22°C. While it has been shown earlier that leaf exudation rates at low temperature do not significantly correlated with broad geographical origin of diverse plant species and phloem loadings types, i.e. symplasmic and apoplasmic (Schrier *et al.*, 2000), the observed significant and homogenous decrease of exudation rates in northern accessions still might indicate a cold acclimation strategy of *Arabidopsis thaliana.* Similar to photosynthetic capacities, cold acclimation of northern accessions might prepare plants more efficiently for a survival of sub-zero temperatures and a significant reduction of sucrose export from leaves into sink tissue might support the accumulation of cryoprotectants and osmolytes (Zuther *et al.*, 2018).

In contrast to southern accessions, correlations between leaf exudation rate, xylem and petiole cross section areas in northern accessions became significantly negative during cold acclimation while xylem and petiole area increased (see Fig. 4). The observed proportional increase of all monitored areas and a similar variance compared to control samples suggests that low temperature did not constrain leaf petiole growth to an extent which was observed for southern accessions which contributes to the phenotypic plasticity of Arabidopsis under low temperature (Atkin *et al.*, 2006). Although sucrose is transported in the phloem and xylem only indirectly contributes to leaf sucrose transport from source to sink tissue, the significance of the cold-induced negative correlation of xylem and petiole areas with sucrose exudation rates might indicate a functional role in cold acclimation of northern accessions. Previously, we found that the northern cold tolerant accession Rsch has a higher growth rate in terms of shoot fresh weight accumulation during cold acclimation than the sensitive accession Cvi (Nagler *et al.*, 2015). Thus, a higher growth rate might also be reflected by increased petiole and xylem areas of northern accessions in the present study. Although it remains speculation at this point, maybe sink strength of northern accessions has been reduced during cold acclimation in order to facilitate carbon and energy supply for maintenance and biosynthesis of protective substances (Demmig-Adams *et al.*, 2018). In parallel to a higher shoot growth rate, this would explain the reduced exudation rates in cold acclimated plants and represent a central trade-off for development under low temperature.

## Acknowledgements

We thank the whole of group of Plant Evolutionary Cell Biology and Plant Development at LMU München for many fruitful discussions and support.

## Author contributions

KS, LA, CN and LS performed experiments, KS and LA performed data analysis. TN analysed data and wrote the manuscript. All authors approved the manuscript.

## Supplementary Information

**Supplementary Figure S1.**
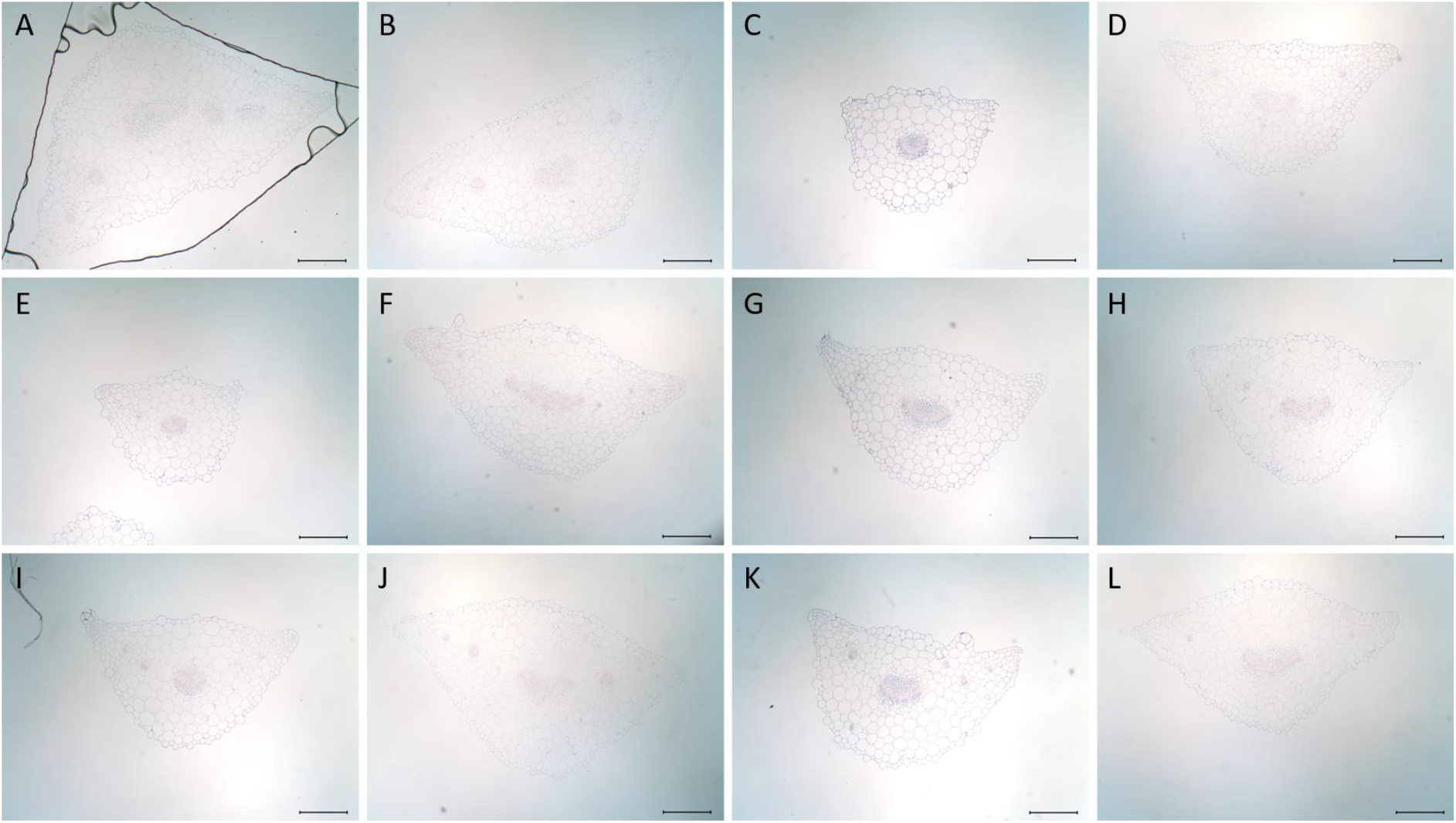
Representative images of leaf base cross sections from different natural accessions of *Arabidopsis thaliana* under control conditions (con) and after cold acclimation (acc). (A) Ct-1 con (B) Ct-1 acc, (C) C24 con, (D) C24 acc, (E) Fei-0 con, (F) Fei-0 acc, (G) Col-0 con, (H) Col-0 acc, (I) Rsch con, (J) Rsch acc, (k) Oy-0 con, (L) Oy-0 acc. Scale bar = 250μm.

**Supplementary Figure S2.**
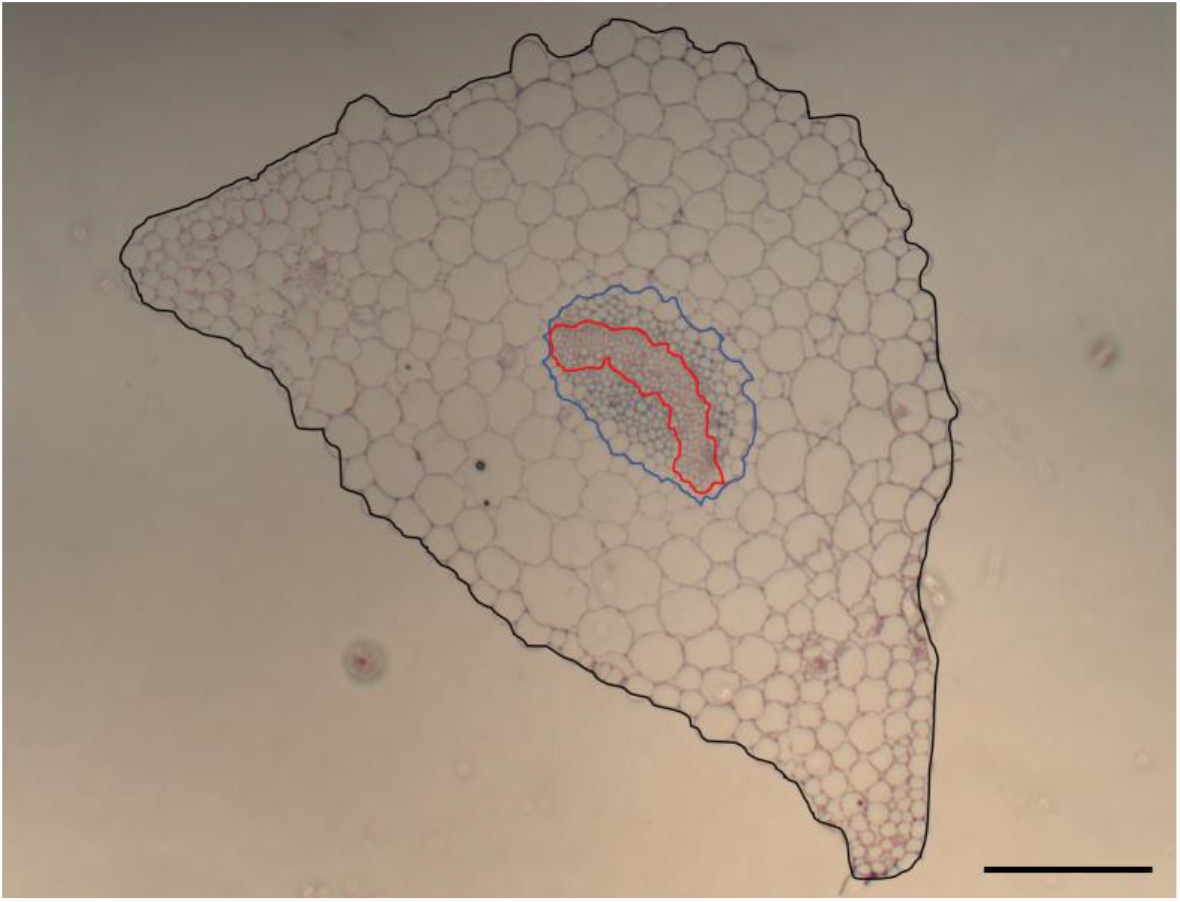
Area definition for petiole area (black line), vascular bundle area (blue line) and xylem area (red). Scale bar = 200μm.

**Supplementary Figure S2.**
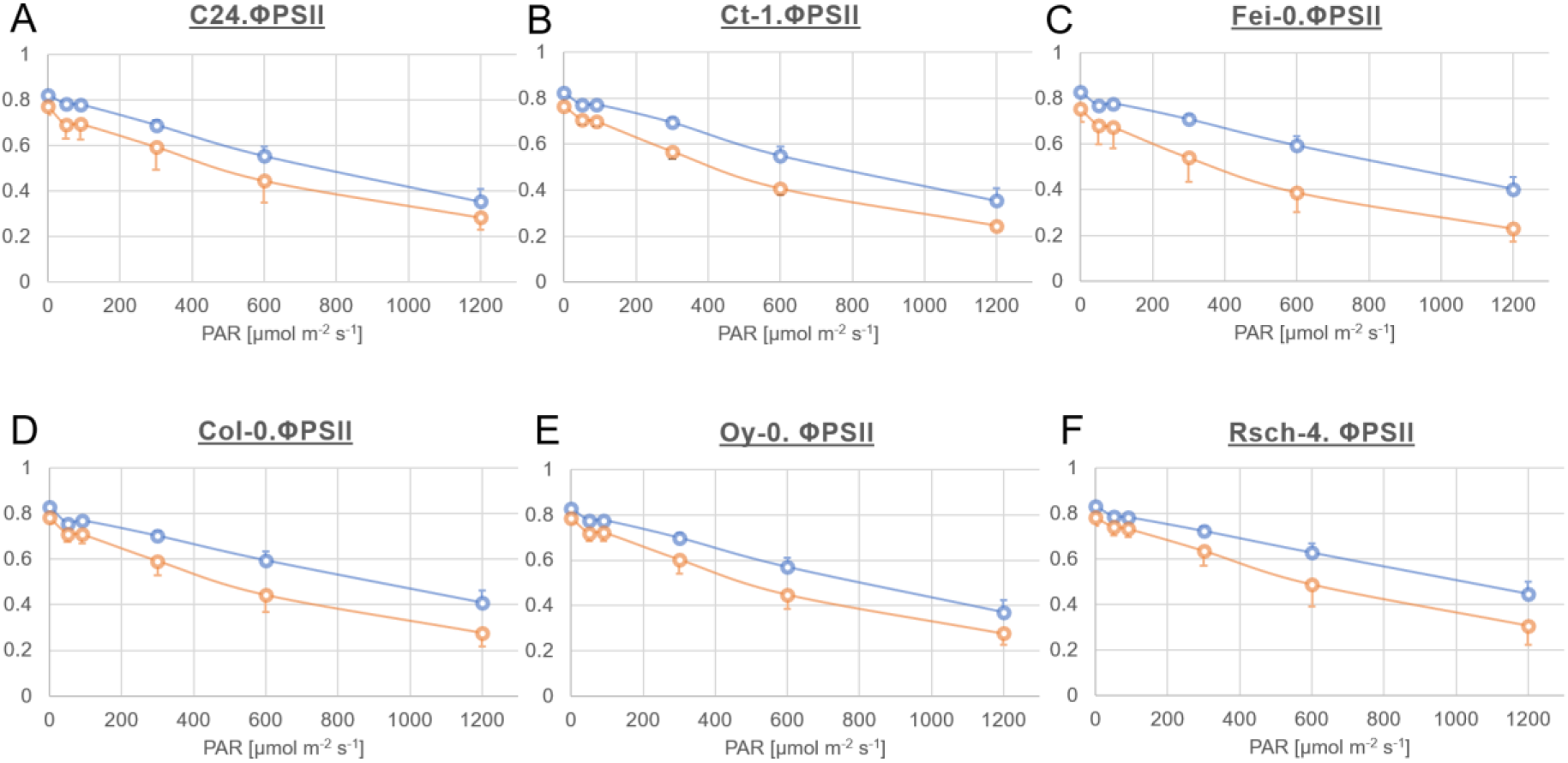
Light response curves of quantum efficiency of PSII (ΦPSII) before and after cold acclimation. Circles and errorbars represent mean values +/− SD (n=3). Blue: control plants; orange: cold acclimated plants (7d, 4°C).

**Supplementary Figure S3.**
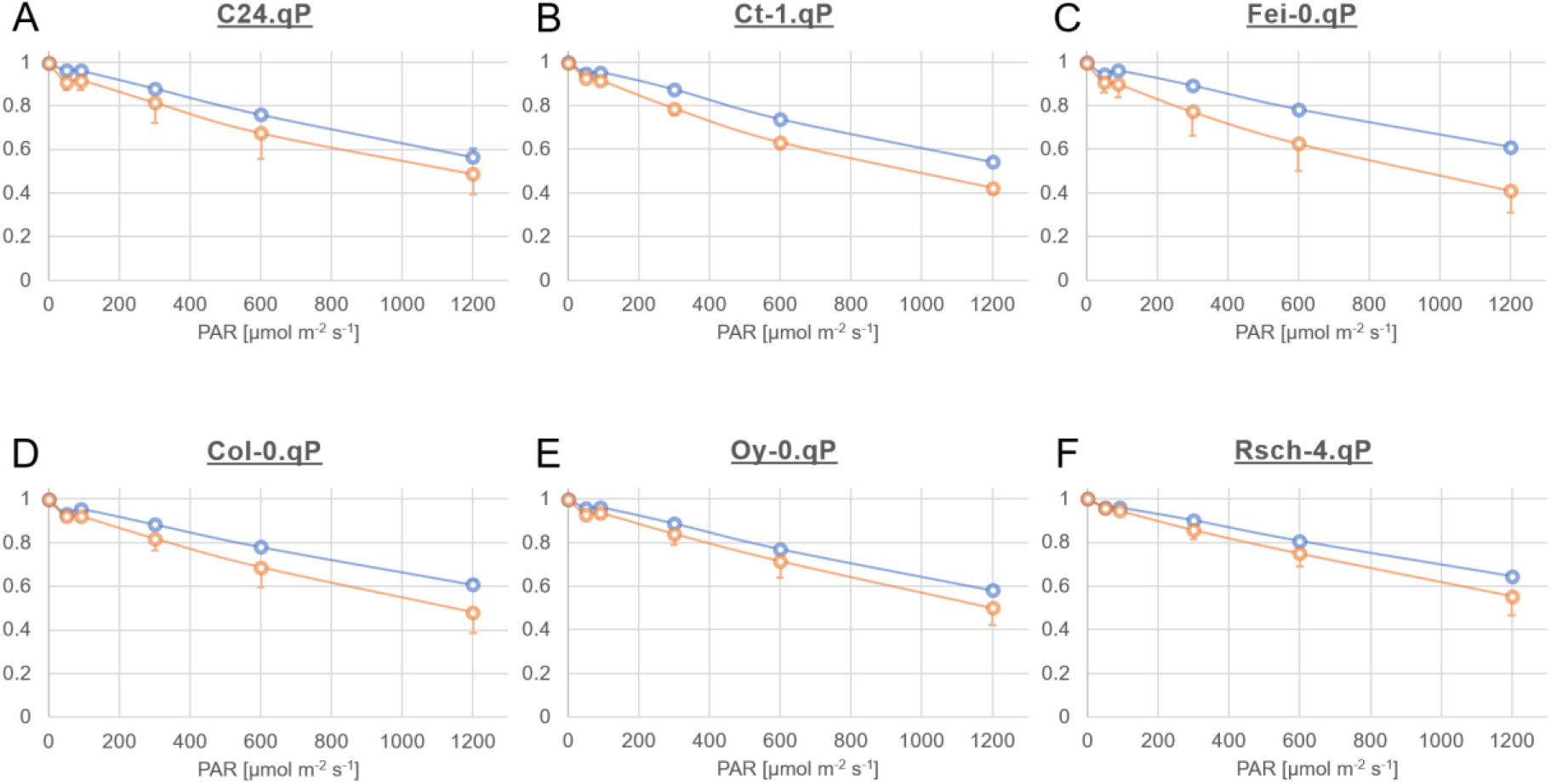
Light response curves of photochemical quenching (qP) before and after cold acclimation. Circles and errorbars represent mean values +/− SD (n=3). Blue: control plants; orange: cold acclimated plants (7d, 4°C).

**Supplementary Figure S4.**
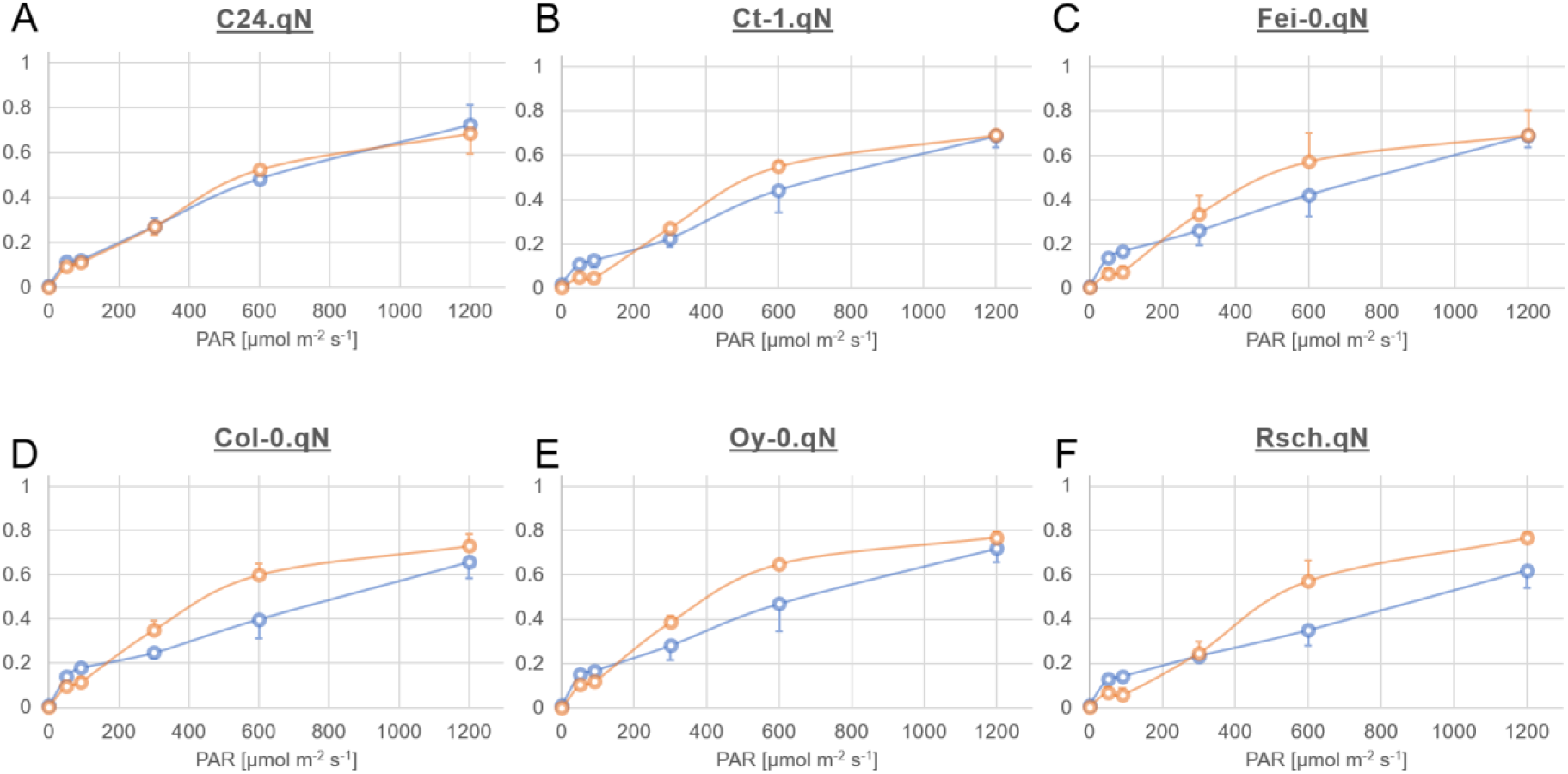
Light response curves of non-photochemical quenching (qN) before and after cold acclimation. Circles and errorbars represent mean values +/− SD (n=3). Blue: control plants; orange: cold acclimated plants (7d, 4°C).

